# First Adaptation of Quinoa in the Bhutanese Mountain Agriculture Systems

**DOI:** 10.1101/691592

**Authors:** Tirtha Bdr Katwal, Didier Bazile

**Affiliations:** Senior Specialist II and National Quinoa Coordinator, Research and Development Center, Yusipang, Department of Agriculture, Ministry of Agriculture and Forests.; Didier Bazile, Senior Researcher in Agrobiodiversity and Agro-ecology at CIRAD, and Quinoa International Focal Point. Considered during the submission my 3 affiliations : CIRAD, UPR GREEN, F-34398 Montpellier, France. GREEN, Univ Montpellier, CIRAD, Montpellier, France. DGDRS, F-34398 Montpellier, France

**Keywords:** Bhutan, Biodiversity, Quinoa, Alternative food crop, Mountain agriculture, Adaptation, Grain yields

## Abstract

Bhutan represents typical mountain agriculture farming systems with unique challenges. The topography, agriculture production systems and environmental constraints are typical of small-scale agricultural subsistence systems related to family farming in the Himalayan Mountains with very low level of mechanization, numerous abiotic stresses influenced by climate and other socio-economic constraints. Quinoa was first introduced in 2015 through FAO’s support to Bhutan as a new crop with the objectives to adapt this versatile crop to the local mountain agriculture conditions as a climate resilient crop for diversifying the farmer’s traditional potato, maize, and based cropping systems, and to enhance the food and nutritional security of the Bhutanese people.

Ten quinoa varieties were evaluated at two different sites representing contrasted mountain agro-ecologies in Bhutan and were tested during the two agricultural campaigns 2016 and 2017. Yusipang (2600 m asl) represents the cool temperate agroecological zone, and Lingmethang (640 m asl) the dry subtropical agroecological zone.The sowing time differed depending on the growing season and elevation of the sites. Results indicate that quinoa can be successfully grown in Bhutan for the two different agro-ecological zones. The grain yields varied from 0.61 to 2.68 t ha^−1^ in the high altitude areas where quinoa was seeded in spring and harvested in autumn season. The grain yield in the lower elevation ranged from 1.59 to 2.98 t ha^−1^ where the crop was sown in autumn and harvested in winter season. Depending on genotypes’ characteristics, agro-ecology and elevation of the sites and variety; crop maturity significantly varied from 92 to 197 days with all genotypes maturing much earlier in the lower elevations where mean minimum and maximum temperatures during the growing season were higher. Quinoa is rapidly promoted across different agro-ecological contexts in the country as a new climate resilient and nutrient dense pseudo cereal to diversify the traditional existing cropping system with some necessary adjustments in sowing time, suitable varieties and crop management practices. To fast track the rapid promotion of this new crop, four varieties have also been released in 2018. In just over three years, the cultivation of quinoa as a new cereal has been demonstrated and partially adapted to the maize-potato traditional cropping systems under the Himalayan mountain agriculture environment. Quinoa is also being adapted to the rice based cropping system and rapidly promoted as an alternative food security crop in the current 12^th^ Five Year national development plan of Bhutan.

## Introduction

Bhutan is a small land locked country with a fragile mountainous environment located in the Himalayan foothills. It is located between China and India with a geographical area of 38,394 square kilometers and a total population of 735,553 persons (PHCB, 2017). It represents a Least Developed Country^3^ with a per capita GDP of US$ 2879.07 (NSB, 2018). Bhutan is mostly mountainous with 70% of its area under forest.

The country is primarily dependent on agriculture and about 62.2% of the population is engaged in subsistence agriculture for their livelihood (GNHC, 2013). Despite agriculture being the primary sector, the country is only able to secure 60% of its cereals, vegetables and animal product needs through domestic production and has to buy over 40% of its food requirement through imports (GNHC, 2013). Bhutanese agriculture represents typical subsistence mountain agriculture where smallholder farmers follow an integrated family farming where agriculture, livestock and forests are intricately linked to meet the household food security. Majority of the farmers grow crops, rear livestock for food, manure and draught power and also depend on forest for fuel, fodder, food, litter and timber. To enhance domestic food production and diversify the farmers existing cropping system, quinoa, a new crop was introduced for the first time to Bhutan in 2015 from Peru (Katwal, 2018; Bazile, Jacobsen and Verniau, 2016) with the support of the Food and Agriculture Organization (FAO) of the United Nations. The rationale for the introduction of quinoa to Bhutan is that FAO has identified quinoa as the most potential crop that can offer food and nutrition security to the world in the next century (Bazile et al, 2015, Jacobsen, 2003, Ruiz et al, 2012).

Lambsquarters (*Chenopodium album*), a quinoa crop wild relative originated from Eurasia, was domesticated in the Himalayan Mountains and it is still possible to find farmers who grow it in India, Nepal, China and Bhutan for its grain or leaves (Partap, T. and Kapoor, P. 1985a and 1985b). It is a historical fact that the use of Chenopodium leaves and seeds for human consumption is not exclusive to the Andean region. A species of Chenopodiaceae was cultivated in the Himalayas a long time ago at altitudes of 1 500–3 000 m asl (Hooker, J.D. 1885, 1952). This species was classified as *Chenopodium album* but also as an unidentified Chenopodium sp., belonged to a complex of two species *C. album* and *C. quinoa* and grown in the Himalayan regions of Punjab. In Bhutan, some farmers continue to grow it and some Chenopodiun sp. as food and vegetables but the identification of species is until today not clear.

In addition, since the eighties, a researcher is growing quinoa on the Tibetan plateau, which presents cold high-desert farming conditions similar to those found on the Altiplano in the Andes. The initial objective was to diversify the mainly barley-based diet of Tibetans by adding vegetable proteins to it through quinoa. The first project was launched in 1984 and the first quinoa plantings took place in 1988. After undergoing training in plant breeding in Mexico and Hawaii, Dr Gongbu Tashi succeeded in adapting quinoa to this mountain region and has now bred local varieties. The area under cultivation is still limited to a few hundred hectares mainly due to difficulties of accessibility. Nevertheless, efforts for adaptation and ongoing work with small farmers have raised yields to nearly two tons per hectare. The painstaking work of plant breeding over different periods through crossing of genetic material from southern Chile, Bolivia and quinoa varieties developed for Mexico has created a new biodiversity of quinoa for high-altitude Himalayan contexts (Gongbu and Wangmu, 1998; Gongbu. 1997).

More recent quinoa trials were conducted in the northern India plains which is quite close to the Himalayan region and have reported that the potential of expanding quinoa cultivation in the Himalayan region is very high (Bhargava et al., 2006; Bhargava1 and Ohri, 2015). Nowadays, several studies (Bazile et al, 2015) have established that quinoa can be cultivated in different growing environment with humidity range of 40 to 90%, at altitudes varying from sea level to 4500 m asl and has the ability to tolerate temperature variation from −8°C to 38°C (PROINPA, 2011, Jacobsen, Jensen and Liu, 2011).

The fundamental objectives of introducing quinoa is to diversify the farmer’s traditional cropping systems through adapting this versatile crop to the Bhutanese subsistence mountain farming systems as a climate resilient crop in order to enhance the food and nutritional security of the Bhutanese people (Katwal, Namgay and Giri, 2018). This paper highlights the results of quinoa demonstration undertaken in 2015, and presents the detail results of replicated trials conducted at two locations in 2016 and 2017.

## Materials and Methods

### Bhutan’s Mountain Agriculture Environment

Before describing the target-growing environment for quinoa adaptation in Bhutan, it is important to provide an overview of the situation of the Bhutanese mountain agriculture where the new quinoa crop is expected to adapt and fit into the existing traditional farming systems. By virtue of being located in the Himalayas, Bhutan is mostly dominated by rugged and steep topography. There is a very large altitudinal variation starting from 100 meters in the south and rising to more than 7,000 m asl in the north. The farming environment is physically challenging with a mountainous terrain. About 5.7% of the total geographical area has a gradient above 100% (45 ° angle), which is prone to very severe soil erosion and soil stability. 43.8% of the area has a slope ranging between 50-100% and 36.5% of the area has slope between 25-50%, while only 14% of geographical area has slope of 0-25% (RNR Statistics, 2015) which indicates the adverse nature of the agriculture land topography. Equally unique is the agro-ecological zones which are categorized into six major groups corresponding with altitude and climatic conditions for agriculture planning and coordination (Table 1). The alpine and the cool temperate zone which is dominated by mountainous terrain has about 53% of the geographical area which clearly indicates the geo-physical setting of the country. Such a fragile geo-physical setting dominated by steep topography makes Bhutan highly vulnerable to any small variation in the weather patterns and the current effects of climate change.

**Table 1.** Major agro-ecological zones of Bhutan

| Agro-ecological zones | Altitude (m.asl) | Temperature (°C) |  | Mean Rainfall (mm) | Proportion of Geographical Area (%) |
| --- | --- | --- | --- | --- | --- |
|  |  | Max | Min |  |  |
| Alpine | 3500-7500 | 12.0 | -1.0 | <650 | 28.6 |
| Cool temperate | 2600-3600 | 22.0 | 1.0 | 650-850 | 23.9 |
| Warm Temperate | 1800-2600 | 26.0 | 1.0 | 650-850 | 18.6 |
| Dry Sub-tropical | 1200-1800 | 29.0 | 3.0 | 850-1200 | 13.1 |
| Humid Sub-tropical | 600-1200 | 33.0 | 5.0 | 1200-1500 | 10.2 |
| Wet Sub-Tropical | 100-600 | 35.0 | 12.0 | 2500-5500 | 5.6 |

Source: Adapted from RNR Research Strategy and Plan Document, MoA, 1992

The cultivated agriculture area is estimated to be only 2.93% of the national geographical area (LCMP, 2010). There are three dominant agriculture land use categories: i-*Chhuzing* (or Wetland) which are terraced paddies for rice cultivation; ii-*Kamzhing* (or dryland) which are rainfed lands; and iii-Horticulture land under orchards and plantations. The predominant crops under orchard and plantations are citrus, apple, arecanut and large cardamom. Among these three land use categories *Kamzhing* is most dominant and constitutes 61.90 % of the agriculture area, *Chhuzhing* covers 27.86% and 10.24% is under orchards and plantations (LCMP, 2010). There are three main distinct cropping systems, which are related to rice, maize and potato based systems with different forms of multiple cropping as one of the simple mechanisms to produce more per unit area (Katwal, 2013).

The dominant cropping systems and crop rotations for different agro-ecological zones are briefly summarized in Table 2. In the *Kamzhing* under the cool and warm temperate zones is potato, wheat or apples based where other crops such as vegetables, mustard, and buckwheat are rotated with cereals or intercropped in orchards. In the dry and humid subtropical areas, maize based cropping systems are predominant with other cereals such as millets and buckwheat, vegetables, legumes and oilseeds are cultivated. Maize + potato intercropping and various forms of multiple cropping are predominant under *Khamzing*. In the terraced wetland or *Chhuzhing* under in the warm temperate zone, farmers mostly grow a single crop of high altitude irrigated rice with some farmers rotating peas, potato, oat and wheat as fodder after rice. The cultivation of a second crop after rice is limited by incidence of early frost and short growing season. In the *Chhuzhing* under wet and humid subtropical areas, rice is followed by mustard, wheat and vegetables in small areas as water is limiting factor after the rice season.

**Table 2.**
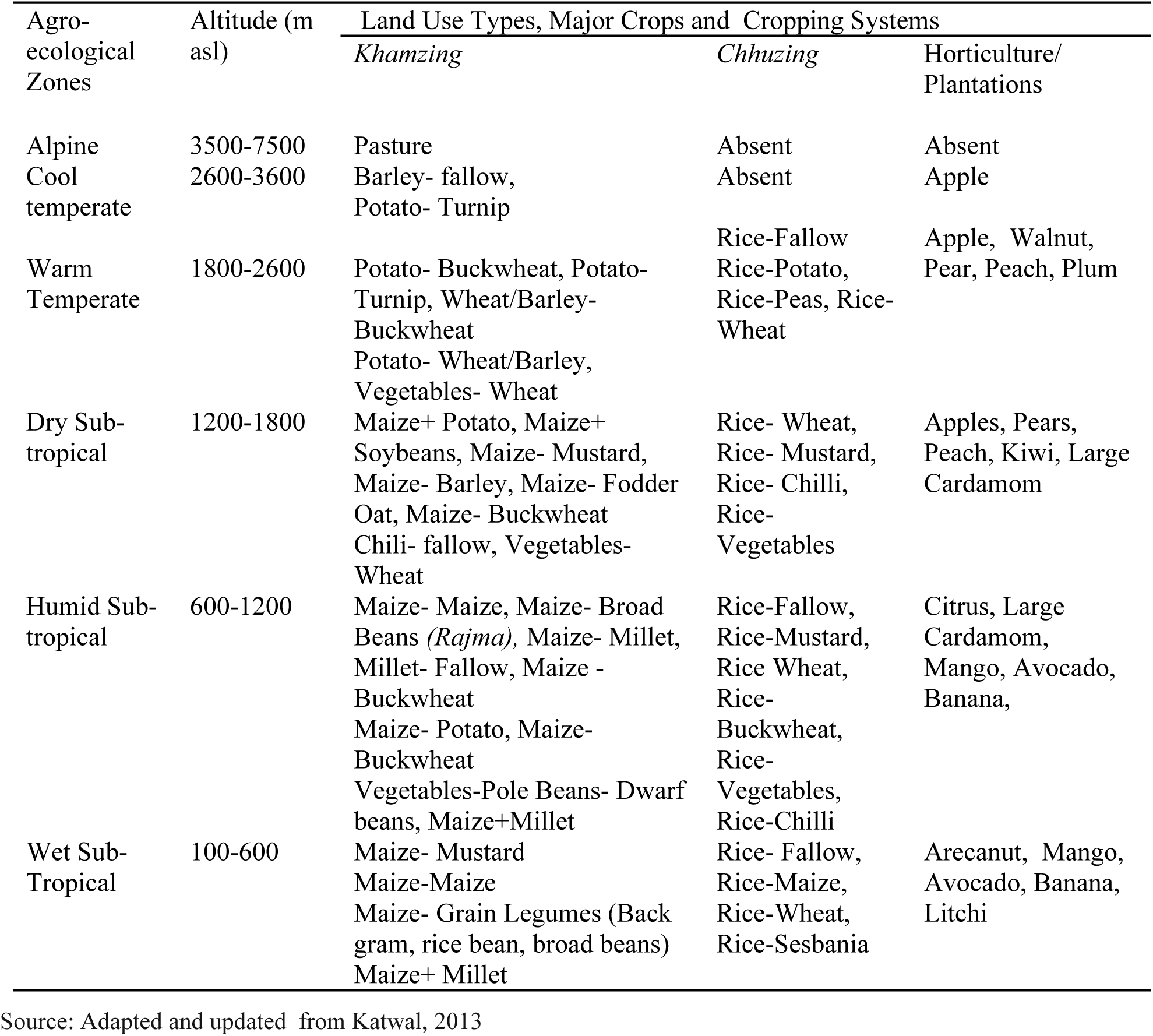
Dominant agriculture land use categories, crops and cropping sequence

| Agro-ecological Zones | Altitude (m asl) | Land Use Types, Major Crops and Cropping Systems |  |  |
| --- | --- | --- | --- | --- |
|  |  | *Khamzing* | *Chhuzing* | Horticulture/ Plantations |
| Alpine | 3500-7500 | Pasture | Absent | Absent |
| Cool temperate | 2600-3600 | Barley- fallow, Potato- Turnip | Absent | Apple |
| Warm Temperate | 1800-2600 | Potato- Buckwheat, Potato-Turnip, Wheat/Barley-Buckwheat<br>Potato- Wheat/Barley, Vegetables- Wheat | Rice-Fallow<br>Rice-Potato, Rice-Peas, Rice-Wheat | Apple, Walnut, Pear, Peach, Plum |
| Dry Sub-tropical | 1200-1800 | Maize+ Potato, Maize+ Soybeans, Maize- Mustard, Maize- Barley, Maize- Fodder Oat, Maize- Buckwheat<br>Chili- fallow, Vegetables- Wheat | Rice- Wheat, Rice- Mustard, Rice- Chilli, Rice- Vegetables | Apples, Pears, Peach, Kiwi, Large Cardamom |
| Humid Sub-tropical | 600-1200 | Maize- Maize, Maize- Broad Beans (*Rajma*), Maize- Millet, Millet- Fallow, Maize - Buckwheat<br>Maize- Potato, Maize- Buckwheat<br>Vegetables-Pole Beans- Dwarf beans, Maize+Millet | Rice-Fallow, Rice-Mustard, Rice Wheat, Rice- Buckwheat, Rice- Vegetables, Rice-Chilli | Citrus, Large Cardamom, Mango, Avocado, Banana, |
| Wet Sub-Tropical | 100-600 | Maize- Mustard<br>Maize-Maize<br>Maize- Grain Legumes (Black gram, rice bean, broad beans)<br>Maize+ Millet | Rice- Fallow, Rice-Maize, Rice-Wheat, Rice-Sesbania | Arecanut, Mango, Avocado, Banana, Litchi |

Source: Adapted and updated from Katwal, 2013

Farm mechanization is highly limited due to steep landscape and as a result, the costs of production for different commodities are very high and problem of erosion and accessibility in marginal areas of mountains. Of the 112,550 hectares (ha) of cultivated area only 32,718 ha is irrigated which means only 29 % of the agriculture area is under irrigation (RNR Statistics, 2015). Thus crop production predominantly depends on seasonal monsoon rains that normally start from late June to September (NEC, 2016) and water for crop production is increasingly becoming scarce and unpredictable due to the increasing variation of precipitation pattern.

Majority of the Bhutanese farmers practices self-sustaining, integrated and subsistence agricultural production system with an average land holding of about three acres where farmers grow a variety of crops under different farming practices and rear livestock to meet their household food security. Owing to the topography dominated by high mountains and deep valleys there is a wide variation of micro climate which requires highly location specific crops and varieties. Many of these crops, varieties and livestock breeds are best adapted to a marginal mountain farming environment for which finding suitable replacements are not straightforward (Katwal et al., 2015).

The average landholding per household is 3.0 acres which is further decreasing due to the rapid rate of land fragmentation that is brought about by the practice of land distribution among children as a right to family land inheritance. Bhutan’s agriculture system is largely traditional with a very minimal use of external inputs like inorganic fertilizers and pesticides. The use of external inputs like chemical fertilizers and plant protection chemicals for agriculture production is estimated to be used by 37% of the farmers in about 19% of the cultivable land, which implies 162,000 acres of cropped area is chemical free (DoA, 2018) making it principally organic by default. Most of the cultivated soils are estimated to have high organic matter content and some soils even exceed 6% organic matter (MOAF, 2011). The National Soil Service Centre estimates that the inorganic fertilizer use in 2016 was 11.9 kg acre^−1^, which is low compared to 13.66 kg acre^−1^ in 2015. In the traditional farming, the application of Farm Yard Manure (FYM) continues to be the major source of plant nutrients which is applied at the rate of 3 to 5 t ha^−1^ (MOAF, 2011).

Bhutan is divided into three distinct climatic zones which are alpine, temperate and subtropical zone. The country has four distinct seasons namely spring, summers, autumn and winter. The spring season which is generally dry starts in early March and lasts until mid-April. The summer season commences in mid-April with occasional showers and continues through the early monsoon rains of late June. The summer monsoon starts from late June through late September that brings heavy rains from the southwest (NEC, 2016). The monsoon brings heavy rains, high humidity, flash floods and landslides, and numerous misty overcast days. The autumn season starts from late September or early October to late November. The dry and cold winter season commences from late November until March. During the winter months occurrence of frost in most part of the country is common and frequent snowfall is experienced at elevation above elevations 3,000 meters.

### Production Challenges of Mountain Agriculture

The most pressing farming constraints of the subsistence Bhutanese farmers are small land holding, dependency on monsoon rains, low farm productivity, high cost of production, low scope of farm mechanization owing to a mountainous terrain, low volume of production and distance from the market. As a result of the steep topography the scope of farm mechanization is highly limited which makes all agriculture operations are labour intensive. The most pressing abiotic challenges are varying precipitation patterns, a short growing season due to the early initiation of frost in the warm temperate and cool temperate areas and increasing climate extremes like drought, hail and windstorms, flash foods and increasing pest problems. Other climate induced abiotic stresses like drought, high temperature, and cold temperature and frost damages to crops decrease farmer’s choices of crops and ability to increasing cropping intensity. A unique and the most pressing problem confronting subsistence Bhutanese farmers is the increasing human wildlife conflicts where crop damage and livestock killings by wildlife are forcing farmers to give up farming. According to the State of Environment Report (NEC, 2016), 55% of the crop damage is attributed to wildlife mainly elephants, wild pigs, deer, monkeys, porcupine and different species of birds. Bhutan has a very strong environment conservation policy which is also one of the four pillars of Gross National Happiness philosophy. The constitution of the Kingdom of Bhutan requires maintaining a minimum of 60% of the area under forest cover. The large proportion of national land area is under forest cover and 51.40 % of the country’s landscape under protected area system that serves as the key habitat for wildlife often attributed for the increasing wildlife impact on agriculture.

Climate change and its impact on subsistence Bhutanese agriculture is an emerging issue where the need for climate resilient crops and climate smart technologies is a priority. To address the increasing impacts of climate induced stresses there is a need to identify and use stress tolerant species which exists but are neglected and underutilized (Ruiz et al., 2013). It has been reported that quinoa has a high degree of resistance to frost and is known to survive a −8°C up to 4 hours depending on the crop stage and variety (Jacobsen, Mujica and Jensen, 2003). Bhutanese mountain agriculture currently faces several hostile production challenges of which most are related to weather and climate. Quinoa has been proven to have exceptional tolerance to hostile environments and is considered a good candidate crop that can help enhance food and nutritional security in the face emerging challenges imposed by climate change (Ruiz et al, 2013). Quinoa is proven for its extreme agro-ecological adaptability and can solve crop adaptation problems in places where climatic and soil conditions are main limiting factors for crop production (Miranda et al, 2012). From its introduction in 2015, quinoa has been included as a priority crop in the current 12^th^ Five Year Plan of the DoA (ARDC, 2018) and rapidly being is evaluated as a climate resilient crop under two different and contrasted agro-ecological zones which have very specific micro environments.

### Trials’ design

First demonstrations of quinoa in Bhutan was conducted in 2015 at Yusipang (2600 m asl), Phobjikha (2900 m asl) and Khangma (2100 m asl) in the research farms with two varieties namely Amarilla Marangani and Amarilla Saccaca. The demonstrations was successful and produced good yields ranging from 2.31 to 2.24 t ha^−1^ (Katwal, Namgay and Giri, 2018). In 2016, 8 more varieties were received from FAO and evaluated in six different locations (Katwal, Namgay and Giri, 2018). Apart from these trials and demonstration two separate quinoa adaptation trials were conducted at Agriculture Research and Development Center (ARDC) research farm at ***Yusipang*** under Thimphu Dzongkhag^4^ and at Agriculture Research and Development Sub-Center (ARDC SC) at ***Lingmethang*** under Mongar Dzongkhag. In both locations, the trials were conducted for two consecutive years in 2016 and 2017. The different quinoa genotypes introduced to Bhutan were evaluated under rainfed dryland. At Yusipang, 10 varieties were evaluated whereas in Lingmethang there were only nine (Tables 4 & 5). Field preparations were done mechanically using powertiller. To prepare the necessary fine seed bed for quinoa; two ploughing was done followed by the pulverization of soil with a rotavator. In both locations about 3 Mt of Farm Yard Manure (FYM) were applied and incorporated in the soil during field preparation. The seed plots were leveled with rakes and 2-3 cm deep furrows were marked uniformly with spade at a spacing 0.50 m. The seeds were uniformly broadcasted in the furrows apparently form lines and covered with a thin layer of soil using a locally made broom.

**Table 3.** Mean Temperature and Precipitation of two trial sites

| Months | Jan | Feb | Mar | Apr | May | Jun | Jul | Aug | Sep | Oct | Nov | Dec |
| --- | --- | --- | --- | --- | --- | --- | --- | --- | --- | --- | --- | --- |
| **Yusipang\* (2600 m asl)** |  |  |  |  |  |  |  |  |  |  |  |  |
| Mean Minimum Temperature °C | -0.9 | 2.1 | 4.4 | 7.5 | 9.8 | 14.2 | 15.4 | 15.3 | 12.5 | 7.3 | 4.7 | 1.3 |
| Mean Maximum Temperature °C | 4.5 | 8.3 | 11.7 | 13.6 | 16.7 | 18.7 | 19.0 | 19.1 | 17.8 | 15.2 | 9.1 | 7.5 |
| Mean Rainfall (mm) | 20.4 | 11.7 | 10.7 | 30.6 | 31.4 | 174.2 | 192.6 | 164.4 | 129.6 | 77.4 | 1.2 | 16.0 |
| **Lingmethang\* (640 m asl)** |  |  |  |  |  |  |  |  |  |  |  |  |
| Mean Minimum Temperature °C | 9.4 | 11.1 | 15.4 | 18.6 | 21.2 | 23.7 | 24.2 | 23.7 | 22.5 | 18.6 | 13.1 | 11.9 |
| Mean Maximum Temperature °C | 23.5 | 23.3 | 26.9 | 30.1 | 32.3 | 32.8 | 33.4 | 32.8 | 32.4 | 30.8 | 27.8 | 26.2 |
| Mean Rainfall (mm) | 8.1 | 23.3 | 23.3 | 57.1 | 103.5 | 143.1 | 138.4 | 134.6 | 104.2 | 46.0 | 9.1 | 1.1 |

\*Data Source for Yusipang: Forestry Research Program, UWICER, Yusipang and RNRRC Annual Reports, 2002-03, 2004-05 and 2007-08
\*\* Data Source for Lingmethang: National Center Hydrology and Meteorology, Ministry of Economic Affairs

**Table 4.**
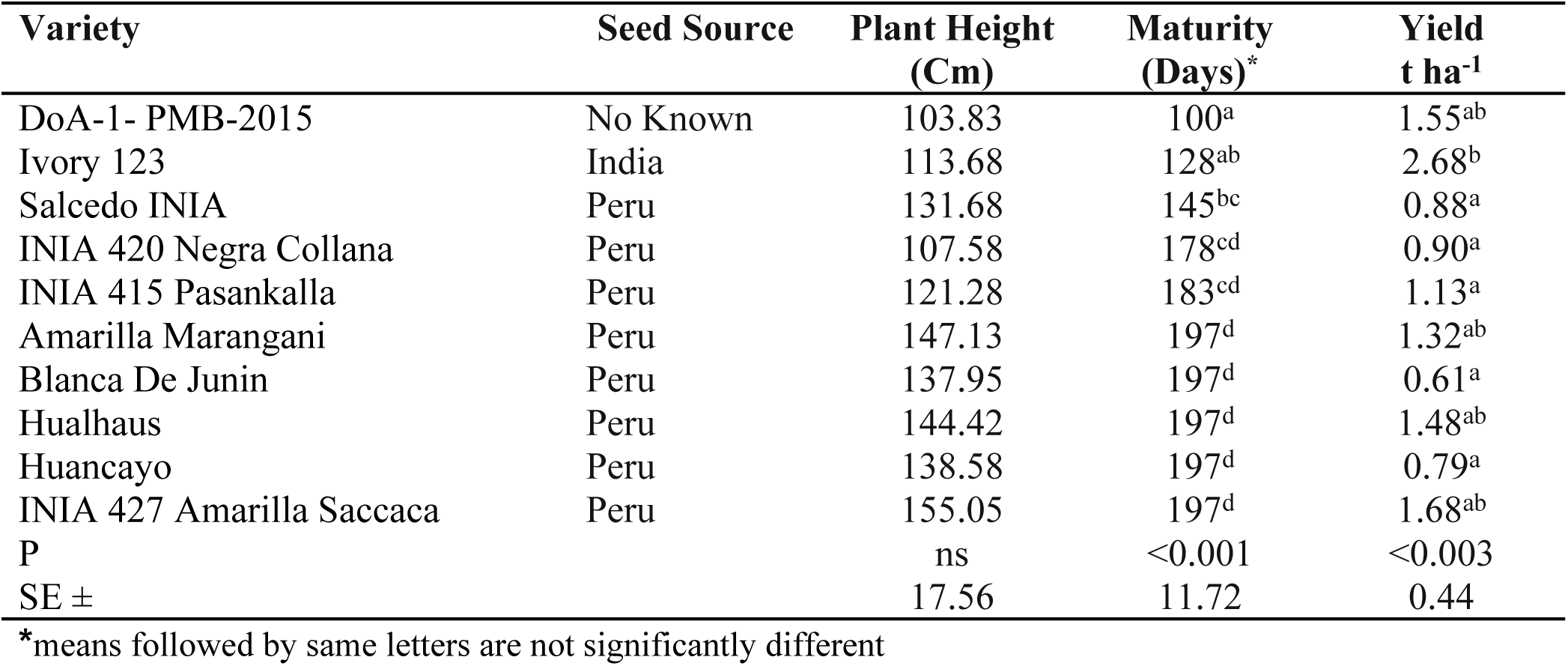
Mean Plant Height, Maturity and Yield of Quinoa Varieties at Yusipang, mean of 2016 and 2017.

**Table 5.** Mean Plant Height, Maturity and Yield of Quinoa Varieties at Lingmethang, mean of 2016 and 2017.

| Variety | Seed Source | Plant Height<br>(Cm) | Maturity<br>(Days) * | Yield<br>t ha<sup>-1</sup> |
| --- | --- | --- | --- | --- |
| INIA 415 Pasankalla | Peru | 106.00 | 92<sup>a</sup> | 1.59<sup>a</sup> |
| Ivory 123 | India | 126.50 | 100<sup>ab</sup> | 2.98<sup>b</sup> |
| INIA 427 Amarilla Saccaca | Peru | 139.67 | 102<sup>abc</sup> | 2.37<sup>ab</sup> |
| Salcedo INIA | Peru | 115.67 | 104<sup>abc</sup> | 1.79<sup>ab</sup> |
| INIA 420 Negra Collana | Peru | 118.53 | 107<sup>abc</sup> | 1.69<sup>a</sup> |
| Amarilla Marangani | Peru | 140.33 | 114<sup>bc</sup> | 2.38<sup>ab</sup> |
| Huancayo | Peru | 134.00 | 114<sup>bc</sup> | 2.51<sup>ab</sup> |
| Blanca De Junin | Peru | 145.57 | 119<sup>c</sup> | 1.95<sup>ab</sup> |
| Hualhaus | Peru | 143.57 | 119<sup>c</sup> | 2.44<sup>ab</sup> |
| P |  | ns | <0.001 | <0.01 |
| SE ± |  | 14.89 | 5.29 | 0.37 |

\*means followed by same letters are not significantly different

In both Yusipang and Lingmethang sites, when the leaves started turning pale yellow, leaf senescence was observed, and grains in the inflorescence became hard on pressing with fingers the crop was considered ready for harvest. The whole plot was harvested manually and the samples from the trials were tied into bundles and dried by hanging in the shade for 10-15 days for curing. The physiological maturity of grains was determined when the seeds from the main panicle were resistant to crushing when pressed between two fingers. The samples were manually threshed and cleaned using local winnowers to obtain the grains for yield estimation. The data from the trials were entered in Microsoft Excel and analyzed with SPSS software. An ANOVA was used to compare the means with level of significance set at 5% (α=0.05).

### Description of the Trial Sites

#### i. Yusipang

At Yusipang the trial was conducted in the research farm at an altitude of 2600 m asl. Yusipang is located at latitude 27°27.5’ to 27°28.2’ N and longitude 89°42.5’ to 89°42.8’ E. Yuispang represents a cool temperate agro-ecological zone and a typical highland mountain farming environment where crop cultivation is limited by cold temperature and frost. The incidence of frost starts by the second fortnight of October and continues till April. Farmers mostly cultivate temperate fruits like apples, peach, plum, walnut and pears. The predominant annual crops cultivated are potato and vegetables mainly the Cole crops. Being a peri-urban area there is a very good opportunity to market any agriculture products and hence the drive to grow fresh vegetables and other agriculture crops is very high.

The predominant soil texture of the site is sandy loams and sandy clay loams (NSSC, 1998). The soils at Yusipang research farm are very acidic to slightly acid, with pH (water) values in the range 4.8 to 6.5. Organic matter levels are low to moderate, with organic carbon contents of about 2.5% and C:N ratios are in the range 10 to 20. Available phosphorous levels are variable but are mostly moderate to high (NSSC, 1998). The mean minimum temperature ranges from −0.9°C in January to 15.4°C in July while the mean maximum temperature varies from 4.5 °C in January to 19.1°C in August. Mean annual rainfall from 2007 to 2017 was 860 mm (Table 3).

At Yusipang, the trial was established on 18^th^ March in 2016 and on 23^rd^ March in 2017. The trial design used was a Randomized Complete Blocks (RCB) with three replications with a plot size of 10 m^2^. Each plot had four rows with a row to row spacing of 0.50 m and the plant to plant spacing was maintained at 0.25 m by thinning. The Indian fertilizers called *Suphala* (16:16:16 NPK) was applied at the rate of 70 kg ha^−1^ and was uniformly incorporated in the furrows drawn with spade before seeding. The seeds were sown uniformly in line and covered with a thin layer of soil using a locally made broom. Weeds were controlled by three hand weeding. The trial was irrigated twice using micro sprinklers before flowering when the crop showed symptoms of moisture stress. Harvesting was done depending on the maturity of the varieties which started in September and was completed by end October for all the 10 varieties.

#### ii. Lingmethang

Lingmethang is situated at 27°15’ 42” N and between 91°10’ 38” and 91°11’ 17” E at an altitude of 640 m asl. This Quinoa trial was conducted at the research farm of the Agriculture Research and Development Sub-Center which is administratively under ARDC Wengkhar. It represents a dry-subtropical agro-ecological zone. The main limiting factors for crop production in the dry-subtropical zone are varying precipitation pattern: rainfed farming that depends on monsoon rains, steep terrain with land slope of 25-50%, and high temperature in summer months especially in valley bottoms. Maize and potato intercropping is very popular. Farmers also cultivate different vegetables, grain legumes, millet, buckwheat, barley and mustard in the dryland.

The mean minimum temperature ranges from −0.9°C in January to 15.4°C in July while the mean maximum temperature varies from 23.4°C in January to 33.4°C in July. The mean annual rainfall from 2007 to 2017 was 792 mm (Table 3). The soils at the Lingmethang are slightly gravelly to extremely gravely and slightly acid to neutral with pH values (water) all above 5.5. The organic carbon and total nitrogen are generally low to moderate and the overall fertility potential and inherent fertility is categorised as slightly poor (NSSC, 2003).

In 2016, the trial at Lingmethang was planted on 20^th^ October and the harvesting dates started from 1^st^ February to 7^th^ March, 2017. The same was trial repeated in 2017 for which the sowing was done on 15^th^ September and the harvesting started from 20^th^ December 2017 and was completed by 7^th^ March 2018. The trial design used was RCB with three replications and the plot size for each variety was 10m^2^ with four rows at a row to row spacing of 0.50 m and plant to plant spacing was maintained at 0.25 m by thinning. Two irrigations with sprinklers of which one was after germination and one at flowering were given. Weed management was done through two hand weeding.

## Results

### Climatic conditions for quinoa adaptation

The climate, cropping season and farming practices of the two trial sites represent a typical mountain environment. The temperatures and precipitation is much lower during the winter season where as the temperature and precipitation during the summer months are higher (Table 3). The winters are much severe in the warm temperate and cool temperate agro-ecological zones (> 1800 m asl) where frost, low precipitation and short growing period is one of the major factors limiting the crop production and choice of crops. In the dry, humid and wet sub-tropical agro-ecological zones the maximum summer temperatures can go above 30°C during April to September which affects seed setting, and higher precipitation causes vivipary in quinoa. Considering the temperature and precipitation that determine the growing season, quinoa was sown from March to April in the warm and cool temperate agro-ecological zones and from September to November in the dry, humid and wet sub-tropical agro-ecological zones. In areas above 1800 m asl represented by Yusipang, quinoa sown from end March to mid April was successful with appreciable yields. The crop sown from September in areas less than 1800 m asl represented by Lingmethang produced good yield and grain quality as the harvest season coincided with dry weather. The soil conditions of both the trials sites was suitable for quinoa with pH values that varied from 4.8 to 6.5 It has been reported that quinoa can tolerant wide range of soil pH from 4.5 to 9 (Pulvento et al., 2010).

### Yusipang study site

At Yusipang (2600 m asl) the statistical analysis of crop maturity (P<0.001) and yield (P<0.003) showed significant differences among the ten varieties evaluated but there was no difference in plant height which is an indication of biomass yield (Table 4). Under the cool temperate agro-ecology (2600-3600 m asl) represented by Yusipang, the days to maturity of 10 varieties ranged from 100 to 197 days (Table 3). The DoA-1-PMB-2015 was the earliest variety which matured in 100 days. Based on the days to maturity the 10 varieties can be grouped into early and late maturing groups. Three varieties maturing in less than 150 days are early while those taking more than150 days are late (Table 4). The yield of the 10 varieties ranged from 0.61 to 2.68 t ha^−1^ (Table 4). The highest yield of 2.68 t ha^−1^was recorded for Ivory 123.

### Lingmethang study site

At Lingmethang (640 m asl) that represents the humid sub-tropical agro-ecological zone, only nine varieties were evaluated for two consecutive years. The statistical analysis of crop maturity (P<0.001) and yield (P<0.01) showed significant differences among the nine varieties but there was no difference in plant height (Table 5). The duration of crop maturity was much shorter at Lingmethang and ranged from 92 to 119 days after sowing (Table 5). The short crop maturity can be explained by the higher mean minimum and maximum temperature (9.4 to 23.5°C) during the growing season from September to February as compared to Yusipang (Table 3). The yield of the nine varieties ranged from 1.59 to 2.98 t ha^−1^ (Table 5) which was much higher as compared to that of Yusipang. The highest yield of 2.98 t ha^−1^ was recorded for Ivory 123.

Quinoa was sown in September as a second crop after maize. The mean maximum temperature at Lingmethang during May to August ranges from 30.1 to 32.4 °C (Table 3) which could affect grain setting. Further, if the crop is sown in February and March, the crop will be exposed to moisture stress in the early growth stages and harvesting will fall during peak monsoon that affects drying, curing and grain quality.

## Discussion

The results from the two study sites representing two types of agro-ecology with different growing environment show that crop maturity and yield vary quite significantly both within and between varieties. The time taken for maturity of all the varieties was much shorter at Lingmethang (92-100 days) as compared to Yusipang (100-197 days), due to the effects of altitude on temperature and affecting the plant development during the agricultural season. The data on crop maturity which is a measure of growing period from seed sowing to harvest from both the sites confirms to the findings that different quinoa genotypes have showed different growing periods (Pulvento et al; 2010). Jacobsen (1997) has reported that under European conditions the growing periods for different quinoa genotypes varied from 109 to 182 days where as in South America the growing period recorded for different genotypes ranged from 110–190 days (Jacobsen and Stølen, 1993). According to Jacobsen (2003), an ideal quinoa variety for grain production should have uniform maturity and early with a growing period of less than 150 days under European northern conditions.

At Lingmethang the total grain yield obtained for all the varieties (1.59 to 2.98 t ha^−1^) was much higher compared to the total grain yield at Yusipang (0.61 to 2.68 t ha^−1^). In the particular case of 2016 year, the means of grain yield (Table 6) of the same 10 varieties were evaluated in six different locations namely Yusipang, Haa, Dawakha, Khangma, Metsham and Trashiyangtse Bhutan representing different agro-ecologies and these varied from 1.22 to 2.57 t ha^−1^ (Katwal, Namgay and Giri, 2018).

**Table 6.** Maturity and mean yield from six location for 10 Quinoa varieties, 2016

| Variety | Days to Maturity* | Grain Yield t ha<sup>-1</sup> |
| --- | --- | --- |
| DoA-1-PMB-2015 | 112.00<sup>a</sup> | 2.52 |
| Ivory 123 | 133.50<sup>ab</sup> | 1.22 |
| Salcedo INIA | 154.17<sup>ab</sup> | 1.24 |
| INIA 420 Negra Collana | 164.83<sup>bc</sup> | 1.33 |
| INIA 415 Pasankalla | 168.50<sup>bc</sup> | 1.51 |
| Hualhuas | 174.00<sup>bc</sup> | 2.39 |
| INIA 427 Amarilla Saccaca | 173.67<sup>bc</sup> | 2.57 |
| Amarilla Maragani | 174.83<sup>bc</sup> | 1.91 |
| Huancayo | 175.50<sup>bc</sup> | 2.23 |
| Blanca de Junin | 187.67<sup>c</sup> | 1.77 |
| **P** | **<0.001** | **ns** |
| **SE ±** | **13.79** |  |

\*means followed by same letters are not significantly different Source: (Katwal, Namgay and Giri, 2018).

In Serbia, Jankovic et al. (2016) reported significantly fluctuation of Quinoa grain yield which ranged from 0.52 to 0.97 t ha^−1^ which was due to soil nutrient contents, and varying levels of precipitation at different crop growth stages. In the Mediterranean climatic conditions in Turkey, experimental grain yields of Quinoa has been found to range from 1.50 to 6.30 t ha^−1^ where as in the farmers field grain yield ranged from 0.50 to 3.50 t ha^−1^ (Yazar et al., 2016). In the Middle East and North African (MENA) countries, it has been reported that average yield of different Quinoa genotypes varied between 1.2 and 1.4 t ha^−1^, while the maximum attainable yield is predicted to go up to 8-10 t ha^−1^ under controlled conditions (Choukr-Allah, et al., 2016; Bazile, et al., 2016).

These trial results and other quinoa demonstrations indicate that quinoa can be successfully promoted under the Bhutanese mountain agriculture environment where temperature and total precipitation in the growing season varies from 4.4-19.1°C and 811mm respectively in the cool temperate agro-ecology. In the humid sub-tropical agro-ecology temperature and total precipitation in the growing season varied from 9.4-32.4 °C and 215mm respectively. This result corroborates the assumptions of Bhargava et al (2006) in India that quinoa cultivation has very high potential to be expanded in the whole Himalayan region. Bhutanese farmers practice three distinct cropping systems which are rice, maize and potato based systems (Katwal, 2013) where quinoa could be easily added for crop diversification. Results from quinoa trials and demonstration conducted in Bhutan in 2015 and 2016/2017 seasons indicated that Quinoa can adapt well and could be grown as an alternative crop in both potato and maize based cropping systems (Katwal, Namgay and Giri, 2018). A study undertaken by Louis et al. (2017) on the challenges and opportunities of climate change for the Bhutanese agriculture sector has also identified quinoa as a potential commodity that is suitable for diversification and expansion of crops in the Bhutanese mountain agriculture systems. This study has also developed quinoa suitability map with areas identified for diversification of quinoa (Figure 2).

**Figure 1:**
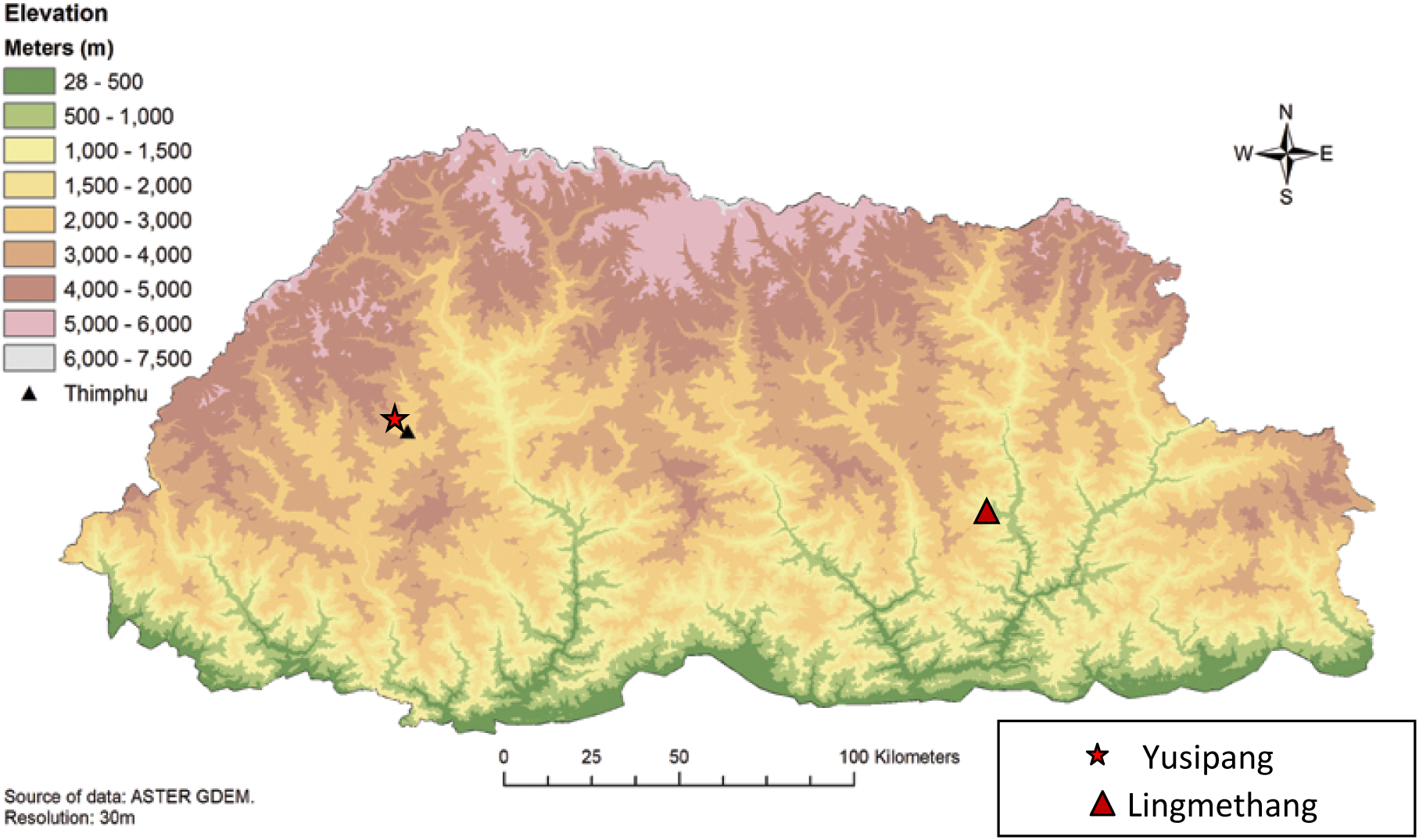
Elevation Map of Bhutan (Source: Parker et al; 2017)

**Figure 2:**
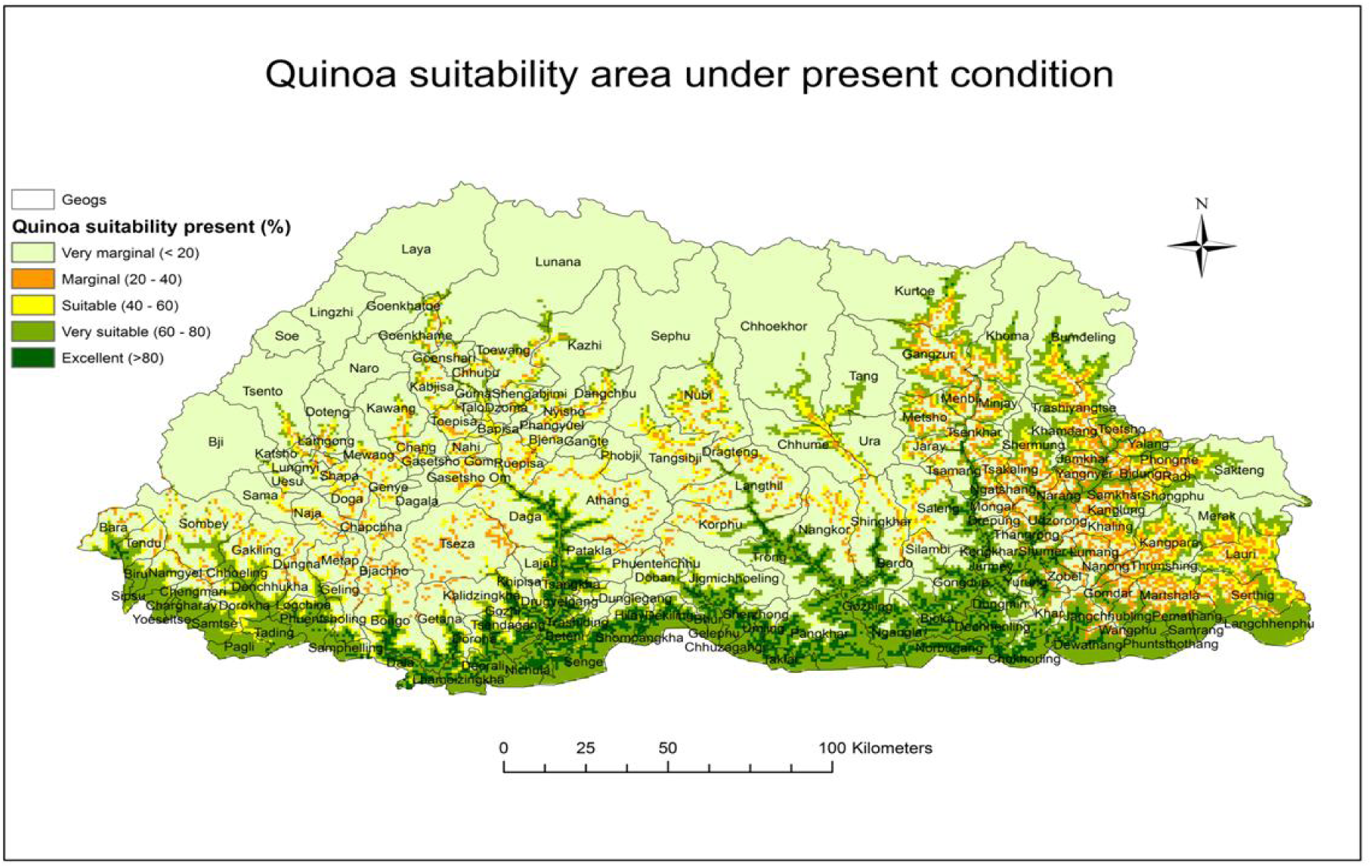
Quinoa suitability area under present condition (Source: Parker et al; 2017)

Quinoa is considered to possess resistance to many adverse abiotic stress like drought, frost and salinity (Jacobsen, Mujica and Jensen; 2003). At Yusipang, the initiation of frost starts as early as the second fortnight of October and continues until the first fortnight of April. The effect of frost damage to quinoa was observed at Yusipang which represent a cool temperate agro-ecology when crop was sown in the first fortnight of March and in August. Some preliminary observation trials on the cultivation of quinoa as a second crop after potato during in the warm and cool temperate areas has indicated that for these areas sowing has to be done by mid of July to escape the frost damage at anthesis. In quinoa crop, the flowering period that commences from first anthesis to the end of flowering is most sensitive to environmental stresses (Bertero and Ruiz, 2008). Further, it has been established that the effect of frost on seed yield of Quinoa can be as high as 51 to 66% when affected at 12 leaf stage and anthesis respectively (Jacobsen, Mujica and Jensen, 2003). It has been reported that despite quinoa’s potential to overcome many abiotic stresses some of the production challenges include sowing under the right conditions to obtain good crop stand from a small seeded crop. After a quick emergence, the management of weed is quite critical in quinoa. The storage and preservation of seed requires special attention because the embryo, with its external position has a short lifespan (Ruiz et al., 2013).

To fast track and rapidly aggressively promote quinoa as a climate resilient crop for food and nutritional security, four varieties have been released for cultivation under different cropping systems and agro-ecology (Table 7). Of the four varieties two are early maturity and two are medium to late in maturity.

**Table 7.** Quinoa varieties released for Bhutanese Cropping System

| Bhutanese Name | Original Name | Origin | Mean Plant Height (cm) | Maturity (Days) | Grain Colour | Mean Yield t ha<sup>-1</sup> |
| --- | --- | --- | --- | --- | --- | --- |
| Ashi Heychum- AM | Amarilla Marangani | Peru | 188 | 173 | Yellow | 1.88 |
| Ashi Heychum- AS | Amarilla Saccaca | Peru | 165 | 170 | Yellow | 2.25 |
| Ashi Heychum- 123 | Ivory 123 | India | 122 | 150 | Brownish | 2.25 |
| Ashi Heychum- TW | DoA-1-PMB-2015 | Unknown | 120 | 140 | Brownish | 1.88 |

Source: Katwal, 2018

## Conclusions

Quinoa, a new crop in Bhutan, has been successfully adapted and acclimatized in just over three years since its first introduction in 2015 to the unique mountain agriculture of this country in the Himalayas. Due to the versatile capacity of the crop to adapt to different growing conditions, quinoa has successfully adapted under the challenging mountain farming environment where different abiotic stresses like varying precipitation patterns, drought, high temperature, frost and cold temperature limit farmer’s choices of food crops and ability to ecological intensification of the cropping systems. The cultivation has been successfully demonstrated to the farmers across the country through on-farm demonstration and four varieties have been recommended with an indicative sowing time for the different agro-ecological zones and cropping systems. In the higher elevations above 1800 m asl spring sowing has been found most feasible. In these areas, a winter crop of quinoa after the harvest of main crops like potato and wheat has been partially successful but more research needs to be done to identify genotypes and critical seeding time so that quinoa can escape frost before anthesis. In the lower elevations below 1800 m asl sowing in autumn in the rainfed dryland where quinoa is cultivated as a rainfed crop after the harvest of potato and maize produced the best results. A winter crop after rice harvest in the terraced paddy fields is currently being evaluated with different genotypes. In this system there is a need to identify suitable varieties with a growing period of less than 150 days, right sowing time and crop management practices.

From a new crop in 2015, quinoa has already spread in all 20 Dzongkhags and across all agro-ecological zones with an estimated area of about 500 acres in 2018. Quinoa is now aggressively being promoted as a climate resilient and a nutrient dense crop across the country in all the agro-ecological zones to enhance food and nutritional security, and as a potential niche organic crop for export market (China & India).

For the Bhutanese farmers, quinoa is a new crop and the overall understanding and management practices of this crop in a mountain farming environment is poor. The technical information available on quinoa is also scanty as the crop has not been evaluated previously in depth in this Himalayan region. Despite a high priority accorded to the promotion of quinoa cultivation an all-out approach of the agriculture research and extension to rapidly upscale the cultivation of quinoa it is faced with numerous challenges. Among them, limited access to quinoa germplasm, absence of best crop management practices under subsistence mountain farming systems, lack of knowledge on pest and diseases, harvesting and processing, package of practices when grown under different agro-ecological conditions and marginal environments are very urgent. Being a new crop the technical knowledge and awareness about quinoa’s nutritional benefits is limited. The absence of marketing channels to sell farmers’ small household surpluses is absent. Further, quinoa has been essentially introduced to enhance the food and nutritional security of the Bhutanese population by promoting it as nutritious health food which requires its inclusion in the traditional cuisines through adequate research and demonstration efforts.

## Acknowledgement

The authors are very grateful to Mr. Padam Lal Giri, Mr. Namgay Wangdi and Mrs. Tashi Dema, Researchers of Agriculture Research and Development Centers at Yusipang and Wengkhar for their support in the implementation of the trials. We are also grateful to Mr. Cheten Thinley, Forestry Research Officer of Ugyen Wangchuk Institute for Conservation of Environment and Research, Yusipang for sharing the climate data for Yusipang. The FAO Bhutan office deserves a very sincere acknowledgement for providing the initial start up seed and budget for quinoa research and development in Bhutan.

## Footnotes

3 “*The Least Developed Countries (LDCs) is a list of developing countries that exhibit the lowest indicators of socioeconomic development, with the lowest Human Development Index ratings of all countries in the world*” according to the United Nations.

4 *Dzongkhag* signifies an “administrative district” in Bhutanese language.

## References

ARDC. 2018. 12^th^ Five Year Plan Strategies for Quinoa Commodity Program. Agriculture Research and Development Center, Yusipang. Department of Agriculture, Ministry of Agriculture and Forest, Thimphu, Bhutan.

Bazile D., Jacobsen S.E., Verniau A. 2016. The global expansion of quinoa: Trends and limits. Frontiers in Plant Science, 7 (622) : 6 p. http://dx.doi.org/10.3389/fpls.2016.00622

Bazile D. (ed.), Bertero H.D. (ed.), Nieto C. (ed.). 2015. State of the art report on quinoa around the world in 2013. Santiago du Chili : FAO, CIRAD, 603 p.http://www.fao.org/quinoa-2013/publications/detail/en/item/278923/icode/?no_mobile=1

Bazile D., Pulvento C., Verniau A., Al-Nusairi M., Ba D., Breidy J., Hassan L., Maarouf I.M., Mambetov O., Otambekova M., Sephavand N.A., Shams A., Souici D., Miri K., Padulosi S. 2016. Worldwide evaluations of quinoa: Preliminary results from post international year of quinoa FAO projects in nine countries. Frontiers in Plant Science, 7 (850) : 18 p. http://dx.doi.org/10.3389/fpls.2016.00850

Bertero, H.D. and Ruiz, R.A. (2008) Determination of seed number in sea level quinoa (*Chenopodium quinoa* Willd.) cultivars. Europ. J. Agronomy 28 (2008) 186–194. Elsevier. doi:10.1016/j.eja.2007.07.002

Bhargava, A. D., Shukla, S., and Ohir, D. (2006). Genetic variability and interrelationship among various morphological and quality traits in quinoa (*Chenopodium quinoa*Willd.) Field Crops Research 101 (2007) 104–116.

Bhargava, A and Deepak Ohri, D. 2015. Quinoa in the Indian subcontinent. In In : Bazile Didier (ed.), Bertero Hector Daniel (ed.), Nieto Carlos (ed.). State of the art report on quinoa around the world in 2013. Santiago du Chili : FAO, p. 511-523.

Choukr-Allah. R., Rao N.K., Hirich A., Shahid M., Alshankiti A., Toderich K., Gill S. and Butt KUR. (2016). Quinoa for Marginal Environments: Toward Future Food and Nutritional Security in MENA and Central Asia Regions. Front. Plant Sci. 7:346. doi:10.3389/fpls.2016.00346.

DoA (2016). Agriculture Statistics 2016. Department of Agriculture. Ministry of Agriculture & Forests. Royal Government of Bhutan. Thimphu, Bhutan.

DoA (2018). Flagship Program on Up-Scaling Organic Sector for Sustainable Socio-Economic Development. Final Draft Proposal. Department of Agriculture, Ministry of Agriculture and Forests, Thimphu, Bhutan.

GNHC (2013). Eleventh Five Year Plan Volume I: Main Document – 2013-18, Gross National Happiness National Happiness Commission, Royal Government of Bhutan. ISBN 978-99936-55-01-5

Hooker, J.D. 1885. The Flora of British India. Volume V. UK, Reeve Kent Publisher.

Hooker, J.D. 1952. Chenopods. Himalayan Journal, 1: 386.

Jacobsen, S.E, Mujica, A. and Jensen, C.R. (2003). The Resistance of Quinoa (Chenopodium quinoa Willd.) to Adverse Abiotic Factors. Food Reviews International. Vol.19 Nos1&2, pp 99–109. DOI: 10.1081.FRI-120018872.

Jacobsen, S.E (2003) The Worldwide Potential for Quinoa (Chenopodium quinoa Willd.), Food Reviews International, 19:1-2, 167–177, doi:10.1081/FRI-120018883

Jacobsen, S. E., 1997: Adaptation of quinoa (Chenopodium quinoa) to northern European agriculture: studies on developmental pattern. Euphytica 96, 41–48.

Jacobsen, S. E., and O. Stølen, 1993: Quinoa – morphology and phenology and prospects for its production as a new crop in Europe. Eur. J. Agron. 2, 19–29.

Jacobsen, S.E., Jensen, C.R and Liu, F. (2011). Improving crop production in the arid Mediterranean climate. Faculty of Life Sciences, University of Copenhagen, Højbakkegård Alle 13, DK-2630 Tåstrup, Denmark.. Field Crops Research 128 (2012) 34–47.ELSEVIER.

Jankovic, S., Popovic, V., Jela Ikanovic3, Rakic, S., Kuzevski, J. and Gavrilovic, M. (2016). Romanian Agricultural Research No. 33, 2016. www.incda-fundulea.ro Print ISSN 1222-4227; Online ISSN 2067-5720

Katwal, T.B. (2018). Quinoa. General Information and Package of Practices, 2018. Field Crops Program, Research and Development Center, Yuispang. Department of Agriculture, Ministry of Agriculture and Forests, Thimphu.

Katwal, T.B. (2013). Popularizing Multiple Cropping Innovations as a Means to Raise Crop Productivity and Farm Income in Bhutan. In Popularizing Multiple Cropping Innovations as a means to Raise Crop Productivity and Farm Income in SAARC Countries. Eds. Musa, M., Azad, A.Kand Gurung, T. (2013). SAARC Agriculture Centre, Dhaka, Bangladesh.

Katwal, T.B, Wangdi, N. and Giri, P.L.(2018). Adaptation of Quinoa in Bhutanese Cropping Systems. Bhutan Journal of Agriculture. Issue II Volume I. In Press. Department of Agriculture, Ministry of Agriculture and Forest.

Katwal, T.B., Dorji, S., Dorji, R., Tshering, L., Ghimiray, M., Chhetri. G.B., Dorji, T.Y. and Tamang, A.M. (2015). Community Perspectives on the On-Farm Diversity of Six Major Cereals and Climate Change in Bhutan. Agriculture 2015, 5, 2–16; doi:10.3390/agriculture5010002. ISSN 2077-0472 www.mdpi.com/journal/agriculture

LCMP (2010). National Soil Service Center and the Policy and Planning Division. Land Cover Assessment Report; National Soil Service Center and the Policy and Planning Division, Ministry of Agriculture and Forest: Thimphu, Bhutan, 2010.

MOAF (2011). National Food Security Paper, Bhutan Climate Summit, 2011. Ministry of Agriculture and Forests, Thimphu, Bhutan.

Miranda, M., Vega-Gálve, A., Quispe-Fuentes. I., Rodríguez, M., Maureira, H. and Martínez, E. A. (2012) Nutritional Aspects of Six Quinoa (Chenopodium quinoa Willd.) Ecotypes from three Geographical Areas of Chile. Chilean Journal of Agricultural Research 72(2) April-June 2012

NEC (2016). Bhutan State of Environment Report. National Environment Commission. Royal Government of Bhutan. Thimphu. ISBN # 978-99936-865-5-2. www.nec.gov.bt

NSB (2018). National Statistical Bureau. Key Indicators. www.nsb.gov.bt. Accessed 23rd November, 2018

NSSC (1998). Technical Report on the Detailed Soil survey of Yusipang RNR Research Centre. Report No. 1(a). Bhutan Soil Survey Project, Research, Extension, and Irrigation Division, Semtokha. Ministry of Agriculture.

NSSC (2003). Technical Report on the Detailed Soil Survey of RNR Research Sub-Centre. Lingmethang, Mongar. Report No: 4.2.02/ SS 18. National Soils Service Center, Semtokha. Ministry of Agriculture.

Parker L; Guerten N; Thi Nguyen T; Rinzin C; Tashi D; Wangchuk D; Bajgai Y; Subedi K; Phuntsho L; Thinley N; Chhogyel N; Gyalmo T; Katwal TB; Zangpo T; Acharya S; Pradhan S; Penjor S. 2017. Climate change impacts in Bhutan: challenges and opportunities for the agricultural sector. Working Paper No. 191. CGIAR Research Program on Climate Change, Agriculture and Food Security (CCAFS). Wageningen, The Netherlands. Available online at: www.ccafs.cgiar.org

Partap, T. & Kapoor, P. 1985a. The Himalayan grain chenopods. I. Distribution and Ethnobotany. Agric. Ecosyst. Environ., 14: 185-199.

Partap, T. & Kapoor, P. 1985b. The Himalayan grain chenopods. II. Comparative morphology. Agric. Ecosyst. Environ., 14: 185-199.

PHCB (2017). Population and Housing Census of Bhutan. National Report. National Statistics. Bureau. Royal Government of Bhutan, Thimphu. www.nsb.gov.btUS

PROINPA, 2011. Quinoa: An ancient crop to contribute to world food security. FAO, Regional Office for Latin America and the Caribbean: Santiago de Chile, 63 p.

Pulvento, C., Riccardi, M., Lavini, A., d’Andria, R., Iafelice. G. and Marconi. E. (2010). Field Trial Evaluation of Two Chenopodium quinoa Genotypes Grown Under Rain-Fed Conditions in a Typical Mediterranean Environment in South Italy. J. Agronomy & Crop Science (2010) ISSN 0931-2250

RNR Statistics (2015). Bhutan Renewable Natural Resources Statistics. RNR Statistical Coordination Section. Policy and Planning Division. Ministry of Agriculture and Forests.

Ruiz K.B., Biondi S., Oses R., Acuña-Rodríguez I.S., Antognoni F., Martinez-Mosqueira E.A., Coulibaly A., Canahua-Murillo A., Pinto M., Zurita A., Bazile D., Jacobsen S.E., Molina Montenegro M. 2014. Quinoa biodiversity and sustainability for food security under climate change. A review. Agronomy for Sustainable Development, 34 (2): p. 349–359. http://dx.doi.org/10.1007/s13593-013-0195-0

Yazar, A. Sezen, S.M., Tekin, S. and İncekaya, Ç. (2016) Quinoa: From Experimentation to Production in Turkey. International Quinoa Conference 2016- Quinoa for Future Food and Nutrition Security in Marginal Environments.

